# Comparing Harmonization Approaches for Protocol-Related Variability in Multisite Diffusion MRI Data

**DOI:** 10.64898/2026.07.07.737018

**Authors:** Kenny Liou, Sophia I. Thomopoulos, Julio E. Villalon-Reina, Hannah Yoo, Yuhan Shuai, Sasha Chehrzadeh, Arvin Arani, Bret Borowski, Robert I. Reid, Prashanthi Vemuri, Clifford R. Jack, Michael W. Weiner, Neda Jahanshad, Paul M. Thompson, Talia M. Nir, Alzheimer’s Disease Neuroimaging Initiative

## Abstract

Diffusion MRI (dMRI) enables assessment of white matter microstructural abnormalities in Alzheimer’s disease (AD), and multisite datasets enable more robust modeling of non-biological variation that can confound analyses. The Alzheimer’s Disease Neuroimaging Initiative (ADNI) includes over 10 dMRI protocols, necessitating robust methods to model protocol-related variability when pooling data. Here, we compared three harmonization approaches: (1) mixed-effects models, (2) ComBat-GAM, and (3) eHarmonize, a reference-based lifespan method. We assessed their ability to reduce protocol-related variability in diffusion tensor imaging fractional anisotropy (FA) and mean diffusivity (MD) while preserving associations with cognitive impairment (CI), and amyloid-beta (Aβ) and tau PET burden in 1,086 ADNI3/4 participants. All approaches yielded more closely aligned FA/MD distributions across protocols. Associations with clinical indicators of CI were highly consistent across approaches, whereas PET associations were less widespread and more variable. Overall, multiple strategies effectively modeled protocol-related variability while preserving AD-related associations.

## I. Introduction

Diffusion MRI (dMRI) is a powerful tool for probing white matter (WM) microstructure inAlzheimer’s disease (AD), where WM differences may provide information on disease-related brain changes [1] that is complementary to volumetric data, PET, and functional neuroimaging. Large multisite studies such as the Alzheimer’s Disease Neuroimaging Initiative (ADNI) [2] enable well-powered analyses of associations between WM integrity and cognitive impairment (CI), amyloid-beta (Aβ) deposition, and tau pathology.

A major challenge in multisite dMRI datasets is substantial variability in scanner hardware, acquisition parameters, and protocol design [3]. These non-biological sources of variation can obscure or inflate biological effects. Because dMRI measures, like diffusion tensor imaging (DTI) fractional anisotropy (FA) and mean diffusivity (MD) are sensitive to differences in acquisition protocols and scanner characteristics, careful modeling of protocol-related variance is needed when data are pooled [5, 6].

Several statistical approaches have been proposed to address non-biological variability in multisite neuroimaging data. Mixed effects models can account for protocol-related variability at the time of analysis. Empirical Bayes approaches, such as ComBat [7] and ComBat-GAM [8], generate harmonized measures while preserving specified biological covariates. Reference-based methods such as eHarmonize [9], align measures to external lifespan reference trajectories. While widely used in neuroimaging, these methods differ in how protocol-related effects are modeled and their impact on AD dMRI analyses remain unclear.

In this study, we compared three harmonization approaches: (1) mixed-effects models, (2) ComBat-GAM, and (3) eHarmonize. We examined their ability to reduce protocol-related variability in FAMD while preserving biologically meaningful associations with AD-related measures: Aβ burden, tau pathology, and of CI.

## II. Methods

### A. Participants and Image Acquisition

Baseline 3T T1-weighted (T1w) MRI and dMRI data were downloaded from the ADNI database. We analyzed dMRI data from 1,086 ADNI3/4 participants: 638 cognitively normal participants (CN), 335 with mild cognitive impairment (MCI), and 113 with dementia (**Table 1**). Participants were scanned with one of twelve ADNI3/4 dMRI protocols (**Table 2**). Key clinical indicators of AD, specifically Clinical Dementia Rating sum-of-boxes (CDR-sob; N=1,036) [10], Aβ-PET centiloids (CLs; N=759) [11], and tau-PET standardized uptake value ratios (SUVRs; N=601) [12], were obtained where available. Cortical Aβ-PET (18F-florbetaben, 18F-florbetapir, 18F-NAV4694) CLs were derived as in [13]. Tau-PET (18F-flortaucipir) burden was defined using the medial temporal SUVR normalized to inferior cerebellar gray matter [14].

**Table 1.**
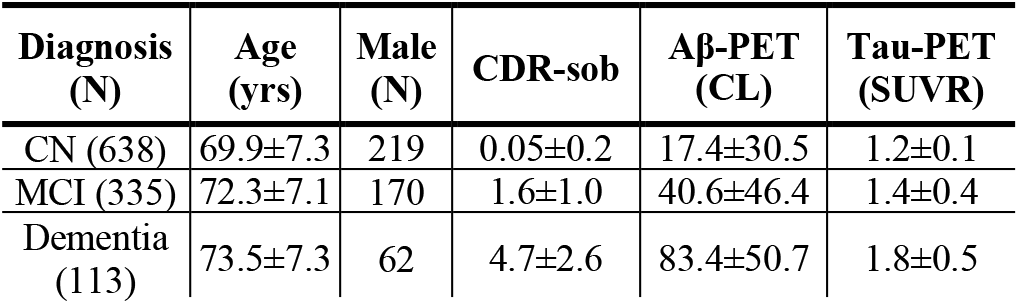
Subject demographics and clinical information.

| Diagnosis (N) | Age (yrs) | Male (N) | CDR-sob | $A\beta$-PET (CL) | Tau-PET (SUVR) |
| --- | --- | --- | --- | --- | --- |
| CN (638) | 69.9±7.3 | 219 | 0.05±0.2 | 17.4±30.5 | 1.2±0.1 |
| MCI (335) | 72.3±7.1 | 170 | 1.6±1.0 | 40.6±46.4 | 1.4±0.4 |
| Dementia (113) | 73.5±7.3 | 62 | 4.7±2.6 | 83.4±50.7 | 1.8±0.5 |

**Table 2.**
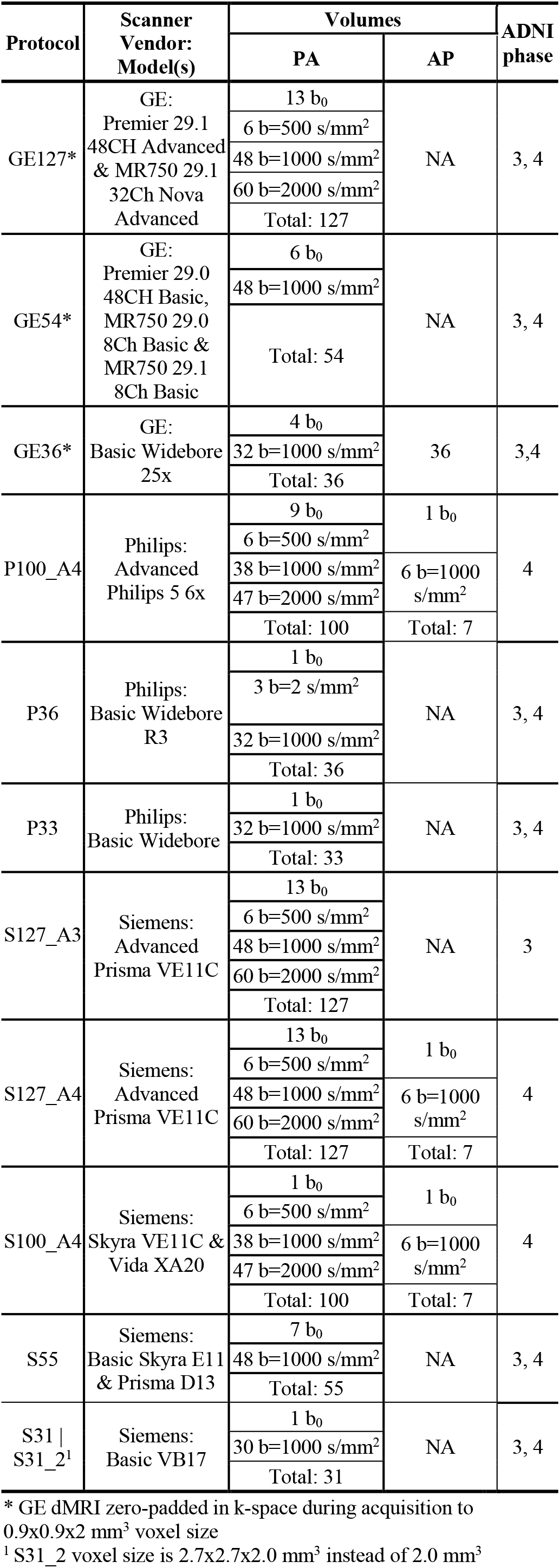
ADNI3/4 dMRI acquisition protocols.

### B. Image Preprocessing and DTI Extraction

All dMRI scans were preprocessed as in [15]. Briefly, scans were denoised and corrected for Gibbs ringing, head motion, and susceptibility-induced distortions. Anterior-posterior and posterior-anterior scans were concatenated when available. Scans were corrected for eddy current-induced distortions and B1 field inhomogeneity. Preprocessed dMRI scans were subsequently warped to each participant’s T1w image. All dMRI and T1w images underwent visual quality control.

DTI FA and MD maps were calculated as in [15]. To ensure comparability across acquisition protocols, tensor fitting included b≤1000 s/mm^2^ volumes only. Using tract-based spatial statistics [16], mean FA and MD measures were extracted from 25 WM regions of interest (ROIs) from the JHU ICBM-DTI-81 [17] atlas, averaged across the left and right hemispheres (**Table 3**).

**Table 3.** Index of 25 JHU WM ROIs.

|  |  |  |  |
| --- | --- | --- | --- |
| **CST** | Corticospinal tract | **SLF** | Sup. longitudinal fasciculus |
| **IC** | Internal capsule | **SFO** | Sup. fronto-occipital fasciculus |
| **ALIC** | Ant. limb of IC | **SS** | Sagittal stratum |
| **PLIC** | Post. limb of IC | **FxST** | Fornix (crus)/Stria terminalis |
| **RLIC** | Retrolenticular part of IC | **UNC** | Uncinate fasciculus |
| **PTR** | Post. thalamic radiation | **TAP** | Tapetum |
| **CR** | Corona radiata | **CC** | Full corpus callosum |
| **ACR** | Ant. CR | **GCC** | Genu of CC |
| **SCR** | Sup. CR | **BCC** | Body of CC |
| **PCR** | Post. CR | **SCC** | Splenium of CC |
| **CGC** | Cingulum (cingulate) | **FullWM** | Full white matter |
| **CGH** | Cingulum (hippocampal) |  |  |
| **EC** | External capsule |  |  |
| **Fx** | Fornix |  |  |

### C. Harmonization of DTI Measures

Four harmonization approaches were applied to regional FA and MD measures across ADNI3/4 dMRI protocols: (1) mixed-effects models, in which protocol was modeled as a random intercept and age, sex, age-by-sex interaction, and ADNI phase were included as fixed effects; (2) ComBat-GAM (All) [8], applied to the full cohort while preserving diagnosis, age, sex, age-by-sex interaction, and ADNI phase; (3) ComBat-GAM (CN) in which only CN participants were used to estimate harmonization parameters while preserving age, sex, age-by-sex interaction, and ADNI phase, with the resulting model subsequently applied to the full cohort; and (4) eHarmonize [9], a reference-based harmonization approach applied separately to each ADNI phase to correct for protocol effects using an external normative dataset.

Two ComBat-GAM implementations were included to evaluate whether the training sample influences downstream findings and highlight a tradeoff: CN-only training provides a more biologically homogeneous reference group but reduces the sample available to estimate protocol effects, whereas full-cohort training increases sample size but may not fully account for biological heterogeneity within diagnostic groups (such as potentially higher variance in disease).

### D. Statistical Comparison of Harmonization Strategies

The harmonization strategies were compared based on (1) their ability to reduce protocol-related variability and (2) their sensitivity to biologically meaningful AD-related associations. To quantify residual protocol-related variability, we tested associations between dMRI protocol and regional FA and MD measures in CN participants before and after applying each strategy. Linear models were fit separately for each ROI and DTI metric, adjusting for age, sex, age-by-sex interaction, and ADNI phase. Protocol was modeled as a fixed effect, and protocol-attributable variance was summarized using partial R^2^. For the mixed-effects strategy, adjusted residuals were derived from models in which dMRI protocol was modeled as a random intercept.

To assess preservation of sensitivity to AD-related associations, regional FA and MD measures were tested for associations with four AD-related metrics: (1) CDR-sob, (2) an MCI versus CN diagnosis, (3) Aβ-PET CLs, and (4) tau-PET SUVRs. For each method, linear models were fit for each ROI and AD outcome, adjusting for age, sex, age-by-sex interaction, and ADNI phase as fixed effects. For Aβ and tau analyses, diagnosis was additionally included as a fixed covariate. For the mixed-effects approach, dMRI acquisition protocol was additionally modeled as a random effect. Association effect sizes are summarized using partial d and partial correlation r for categorical and continuous AD indicators, respectively. The false discovery rate (FDR) procedure [18] was used to correct for multiple comparisons across 25 ROIs.

## III. Results

### A. Removal of Protocol Effects After Harmonization

As an illustrative example, unharmonized FullWM FA and MD measures in CN participants showed substantial protocol-related differences across acquisition protocols (Variance explained by protocol: partial *R*^*2*^≥0.24, *P*<0.001; **Fig. 1A**). All four harmonization approaches reduced these differences, resulting in more closely aligned distributions across protocols (**Fig. 1B-E**). Across regional FA and MD measures, residual protocol-attributable variance was low after harmonization (partial *R*^*2*^<0.03), with no significant residual protocol associations (*P*≥0.11).

**Figure 1.**
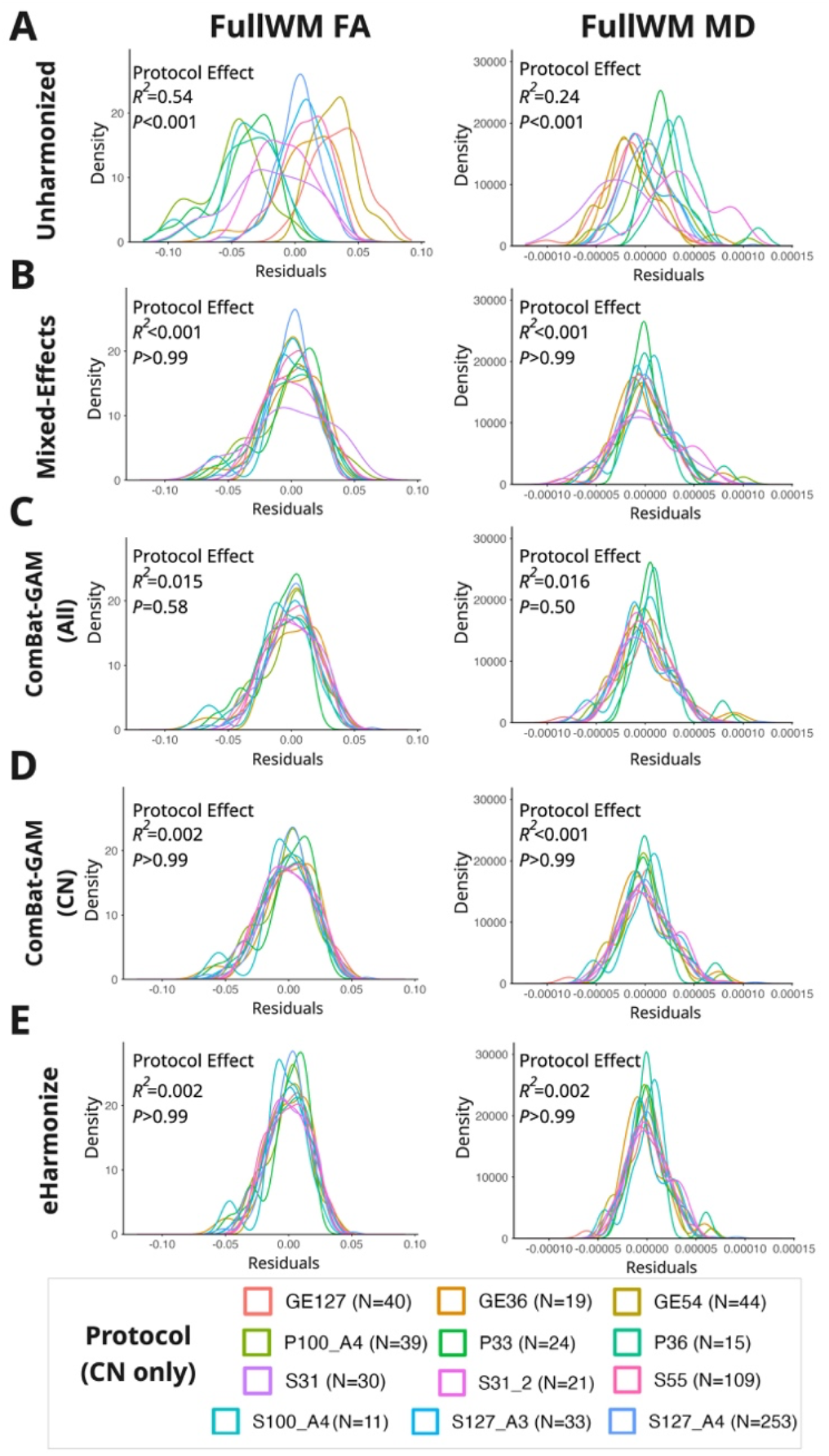
Density plots of unharmonized and harmonized FullWM FA and MD residuals in CN participants show reduced protocol-related differences after harmonization. To minimize the influence of demographic differences on observed variability, FA and MD were adjusted for age, sex, age-by-sex interaction, and ADNI phase. For the mixed-effects approach, dMRI protocol was also included as a random effect. Separate fixed-effects analyses quantified residual protocol-related variance.

### B. AD-related Associations Across Harmonization Strategies

Across strategies, lower FA and higher MD were widely associated with greater CI (i.e., greater CDR-sob and an MCI diagnosis; **Fig. 2A, B**). Strongest effects were localized to regions including the CC, CGH, and Fx. In contrast, Aβ and tau associations were less widespread. Greater Aβ burden was associated with higher FA (e.g., ALIC, SFO) and higher MD (CGH) (**Fig. 2C**). Greater tau burden showed directionally similar associations to CI, with lower FA and higher MD in a smaller number of ROIs (**Fig. 2D**). The number of significant ROI associations was highly consistent for CI but varied more for the weaker PET associations, particularly tau-PET MD (**Table 4**).

**Figure 2.**
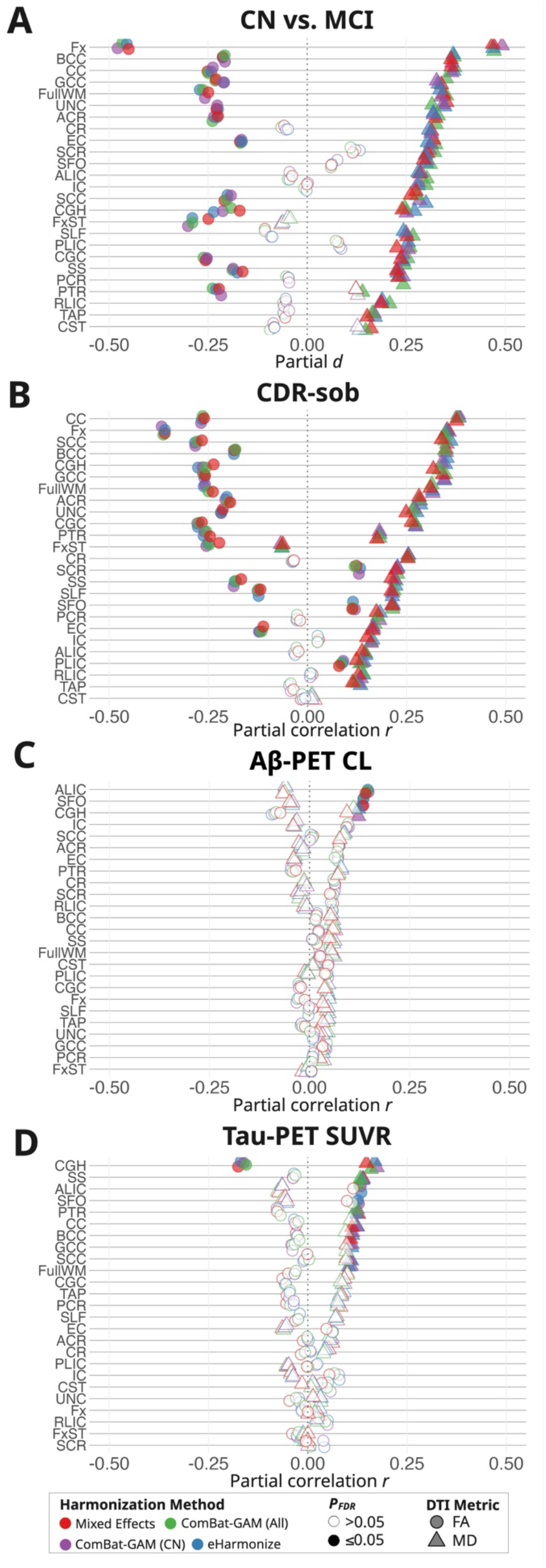
Effect sizes for regional FA and MD associations with (A) MCI diagnosis, (B) CDR-sob, (C) Aβ-PET CLs, and (D) tau-PET SUVRs after application of each harmonization method.

**Table 4.**
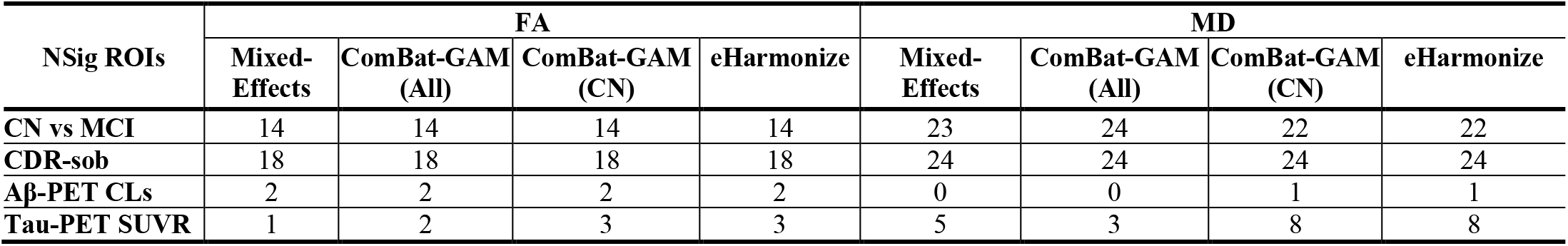
Number of significant FA and MD ROI associations across AD-related metrics and harmonization approaches (P_FDR_≤0.05).

## IV. Discussion

Overall, all harmonization approaches lowered protocol-related variability in FA and MD measures, supporting the effectiveness of harmonization in mitigating inter-site differences that are well documented in diffusion MRI studies [7, 8, 9]. All strategies yielded highly consistent associations between regional FA/MD measures and clinical indicators of CI, including MCI diagnosis and CDR-sob. Associations with Aβ-PET and tau-PET burden were less widespread and showed greater variation across harmonization methods, particularly for tau-PET MD. However, these differences may partly reflect the smaller PET subsamples; we also lack a known ground truth against which preservation of biological signal could be directly evaluated. These findings suggest that the choice of harmonization strategy may have limited influence on stronger clinical associations, but weaker biomarker associations may be more sensitive to the choice of harmonization approach.

Rather than identifying a universally optimal method, our findings highlight the distinct strengths and use cases of each strategy. Mixed-effects models account for protocol-related variability at the time of statistical analysis and may be preferred when a transformed dataset is not required. In contrast, ComBat-GAM and eHarmonize generate harmonized measures for downstream analyses. ComBat-GAM can flexibly model nonlinear covariate effects, while eHarmonize aligns measures to external lifespan trajectories, facilitating harmonization to a common reference and application to new datasets. These practical distinctions are summarized in **Table 5**.

**Table 5.**
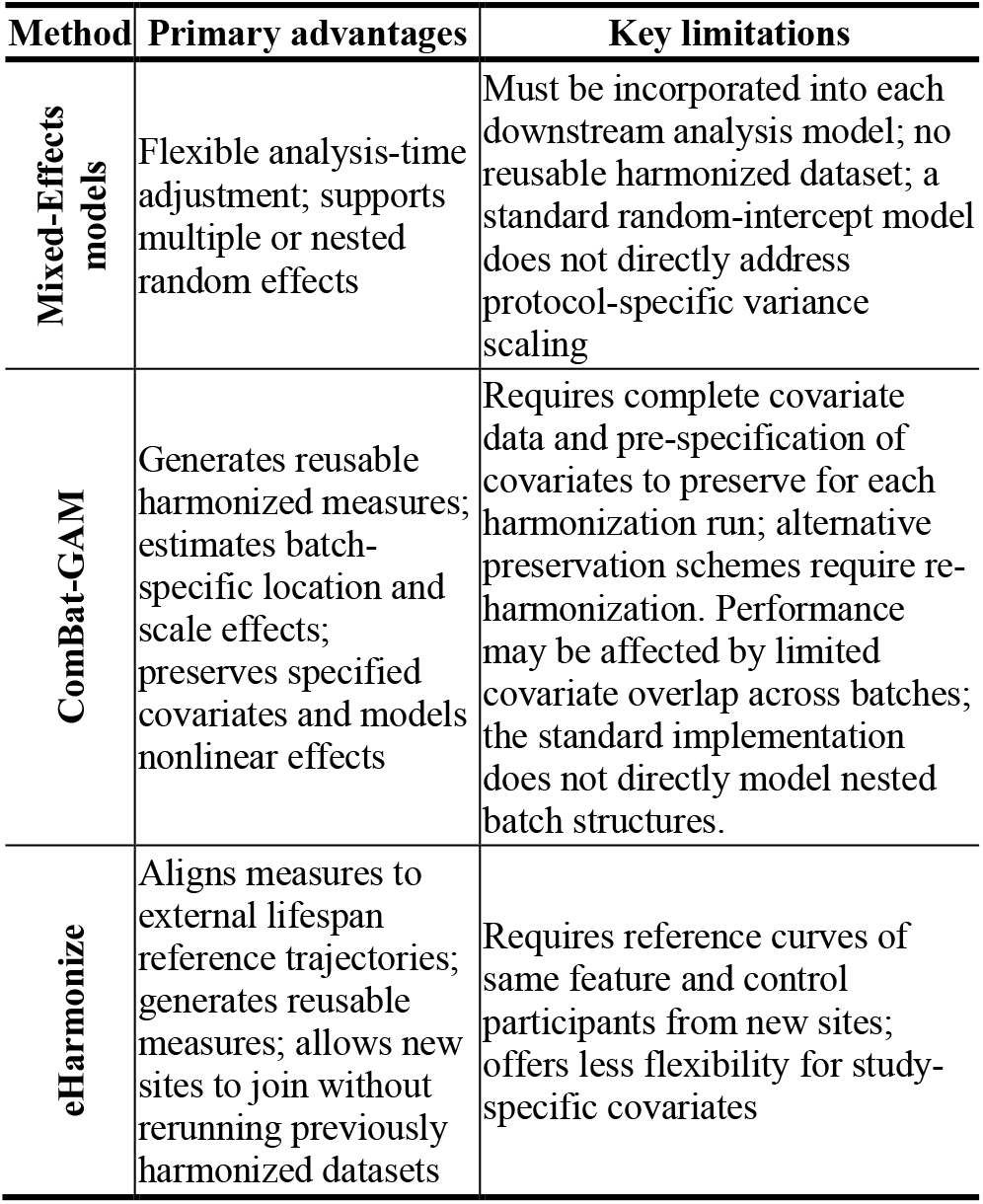
Advantages and limitations of harmonization strategies.

Future work should test these strategies in more heterogeneous cohorts, evaluate additional diffusion metrics and downstream modeling tasks, and examine frameworks that address higher-order distributional differences such as skew and kurtosis. These extensions will help define the circumstances in which each strategy is most appropriate for multisite dMRI studies.

## Acknowledgments

This work was supported by AARG-23-1149996, R01AG058854, R01AG087513, RF1AG057892, RF1NS136995, U19AG024904, U01AG068057, S10OD032285.

